# Integrating Spatially Adjusted Protein Summaries for Survival Prediction in Spatial Proteomics

**DOI:** 10.64898/2026.06.08.730964

**Authors:** Seungjun Ahn, Eun Jeong Oh, Diddier Prada, Ali Shojaie

## Abstract

Recent advances in spatial proteomics, particularly imaging mass cytometry, enable the measurement of protein expression at the single-cell level while preserving a spatial context. Conventional survival analyses, however, typically rely on patient-level averages of protein intensities and therefore overlook spatial heterogeneity and tissue architecture. To address this limitation, we introduce a framework that incorporates spatial information into survival modeling by generating spatially adjusted protein summaries (SAPS). In this approach, cell-level protein intensities within each patient are modeled using spatial spline regression to capture spatial trends. From these models, we extract two complementary features: a spatially adjusted mean expression and a residual variance that reflects cell-to-cell variability unexplained by spatial effects. These summaries are then incorporated into Cox proportional hazards models in combination with clinical covariates. We further show that our estimator is asymptotically equivalent to an oracle estimator under mild regularity conditions. In simulation studies, our proposed framework achieved improved predictive performance compared to other alternative methods. The application of the method to breast cancer imaging mass cytometry data indicate that spatially adjusted summaries may enhance survival prediction and reveal biologically interpretable spatial protein patterns, suggesting high translational potential. This methodology offers an efficient means of translating complex spatial proteomics data into patient-level features, providing both improved survival prediction and new insights into the role of spatial heterogeneity in cancer outcomes. R package is available on the Comprehensive R Archive Network repository at https://cran.r-project.org/web/packages/SurvSPro/index.html

## 1 Introduction

Recent advances in spatial proteomics (SP, hereafter) imaging techniques, such as imaging mass cytometry (IMC) (Chang et al., 2017) and multiplexed ion beam imaging (Schapiro et al., 2017), have enabled simultaneous quantification of protein expression while preserving spatial information at single-cell resolution. As a result, many spatial proteomics datasets have become publicly available, allowing researchers to investigate the functional roles of spatially organized cells and interactions across different spatial regions. (Wang et al., 2025). Recent studies have shown that the tumor microenvironment (TME), a complex ecosystem composed of immune cells, non-immune cells, stromal components, and blood vessels, plays a key role in cancer progression (Quail & Joyce, 2013). TME characteristics have also been used for cancer detection (Binnewies et al., 2018) and for predicting response to treatments such as immunotherapy (Murciano-Goroff et al., 2020). SP data are particularly useful in this context because the spatial distribution of immune cells within the TME is critical for predicting patient outcomes and evaluating immunotherapy response (Benimam et al., 2025; Wang et al., 2023). We focus on breast cancer (BC), which caused an estimated 2.3 million new cases and approximately 666,000 deaths in 2022 (Bray et al., 2024). BC tissues provide a relevant setting for spatial modeling because tumor architecture often includes irregular invasive fronts and variable stromal density (Bahcecioglu et al., 2020).

Time-to-event outcomes (e.g., overall survival) are commonly analyzed as primary endpoints in cancer research. A Cox proportional hazards model (Cox, 1972) is typically used to regress multiple covariates associated with patient survival. In the context of SP data, previous studies have incorporated features such as cell-type fractions (Aung et al., 2025) or protein expression summaries (Yaghoubi Naei et al., 2024) as covariates in Cox models. However, these approaches often ignore the spatial organization of cells, which may limit the ability to identify prognostic patterns arising from cell-cell spatial interactions. A recent study proposed a Cox-model-based framework (TopKAT) that incorporates spatial information derived from the spatial organization of cells in SP data (Samorodnitsky et al., 2026). However, TopKAT primarily operates on cell-type labels and locations to extract topological structures without leveraging the underlying molecular intensity information within cells, limiting its ability to detect spatially varying functional signals.

To address this limitation, we develop a statistical method that incorporates spatial information into survival modeling through spatially informed protein summaries. Our approach consists of two steps. First, cell-level protein intensities within each patient are modeled using generalized additive model (GAMs), in particular, spatial spline regression to capture spatial trends across the tissue. From these models, we derive two complementary summaries for each protein marker, a spatially adjusted mean expression and a residual variance that reflects cell-to-cell variability unexplained by spatial effects. Second, these summaries, which capture different aspects of spatial protein expression patterns, are incorporated into Cox models together with clinical covariates to assess their association with patient survival. A major advantage of the proposed method is that it preserves spatial information in protein expression while summarizing cell-level data at the patient level, allowing the effects of spatially informed protein summaries on patient survival to be estimated using hazard ratios within the standard Cox regression framework.

The remainder of this article is organized as follows. Section 1 provides background and motivations. Section 2 introduces our proposed methodology for spatial proteomics with survival data. In Section 3, we conduct simulation experiments, comparing our method with other existing methods. In Section 4, we apply our proposed method to an IMC dataset of BC patients. Lastly, discussions and future directions are provided in Section 5.

## 2 Methods

### 2.1 Spatial Spline Modeling and Summaries

We consider single-cell spatial expression data for a single protein marker of interest, where each cell has a measured intensity value for that protein and a pair of spatial coordinates. For example, in our real data application, the marker of interest is CD3 epsilon (CD3E). For patient *i* = 1, …, *N*, let *J*_*i*_ denote the set of that patient’s tumor cells, where |*J*_*i*_| = *m*_*i*_ is the total number of tumor cells for patient *i*; if tumor labels are not available, we include all cells.

We consider single-cell spatial expression data where each cell has a measured protein intensity and a pair of spatial coordinates. For patient *i* = 1, …, *N*, let *J*_*i*_ denote the set of that patient’s tumor cells, where |*J*_*i*_| = *m*_*i*_ is the total number of tumor cells for patient *i*; if tumor labels are not available, we include all cells. For each cell *j* ∈ *J*_*i*_, let *y*_*ij*_ be the protein intensity with its corresponding spatial location (*u*_*ij*_, *v*_*ij*_).

To capture smooth spatial trends in protein expression, we model the intensity using a two-dimensional thin-plate spline,

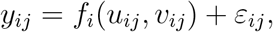

where E[*ε*_*ij*_] = 0 and *f*_*i*_(*·*) is estimated using a generalized additive model (GAM), similar to Vasconcelos et al. (2025), with a thin-plate basis and basis dimension *k* (Wood, 2017). Thin-plate splines provide a flexible, non-parametric way to represent spatial variation, allowing complex tissue structures to be modeled without imposing strong assumptions about local patterns. Before fitting the model, the spatial coordinates are rescaled to the interval [0, 1] to improve numerical stability.

Let 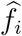 denote the fitted value and define residuals as 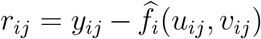. We construct a polyon *A*_*i*_ from the labeled cell coordinates by taking the convex hull of those points. Then, we summarize each patient’s spatial profile using two quantities. First, the spatially adjusted mean is obtained by the tumor area average of the fitted surface over *A*_*i*_,

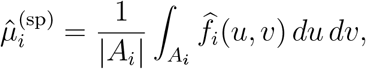

which we approximate by averaging predictions on a uniform grid that evenly covers *A*_*i*_. This reflects the expected expression at a randomly chosen location within the tissue, which is useful in cross patient comparisons, prognostic modeling, and reporting tissue level summaries that are not confounded by variable cell density or sampling design. Second, we quantify local heterogeneity with the residual variance

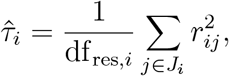

where df_res,*i*_ is the residual degrees of freedom returned by the fitted model for patient *i*. Larger values of *τ*_*i*_ indicate more spatially heterogeneous expression, which may signal disordered or complex tissue architecture.

### 2.2 Survival Modeling

We observe data (*T, δ*, ***X***) from *n* patients, where *T* is the observed time, *δ* is the censoring indicator, and 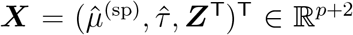 represents the final covariate vector, which consists of two spatially adjusted summaries of the protein introduced in the previous section, along with *p*-dimensional demographic and clinical covariates (i.e., molecular subtype indicators and other clinical characteristics, such as ER/HER2 status, stage, grade, or treatment history). In the proportional hazards model, also known as the Cox model (Cox, 1972; Cox & Oakes, 1984), the hazard function *h*(*t*) is specified as

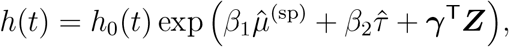

where *h*_0_(*t*) is an unspecified, non-negative baseline hazard function and ***θ*** = (*β*_1_, *β*_2_, ***γ***^T^)^T^ ∈ ℝ^*p*+2^ is the parameter vector. The corresponding Cox partial likelihood is

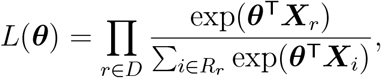

where *D* is the index set of observed event times and *R*_*r*_ is the risk set consisting of all patients at risk at time *t*_*r*_. Herein, *β*_1_, *β*_2_, and ***γ*** denote the regression coefficients associated with the spatially adjusted mean intensity, residual spatial variance, and clinical covariates, respectively. The parameters ***θ*** are estimated by maximizing the Cox partial likelihood using the Efron approximation for tied event times (Efron, 1977), implemented via the `coxph` function in the R package `survival` (Therneau & Grambsch, 2000). For patient *i*, the fitted model yields an estimated risk score

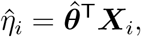

where larger values of 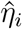 indicate higher risk and poorer expected survival outcomes.

### 2.3 Asymptotics of the two-stage procedure

Let 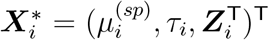 denote the true counterpart of the covariate vector ***X***_*i*_ defined in Section 2.2, where the first-stage estimates 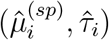 are replaced by their population values, and let 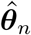 maximize the Cox partial likelihood based on ***X***_1_, …, ***X***_*n*_. Denote *m* = min_*i*_ *m*_*i*_, where *m*_*i*_ is the number of cells for patient *i*, and 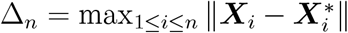. Throughout, we assume the following regularity conditions:

(A1) Δ_*n*_ = *O*_*p*_(*δ*_*n,m*_) for a deterministic sequence *δ*_*n,m*_ that collects the uniform first-stage errors, namely the spline estimation error 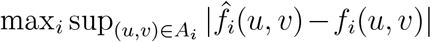, the quadrature error in approximating the area average over *A*_*i*_, and the residual-variance estimation error max*_i_*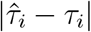.

(A2) 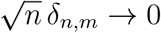.

(A3) The covariate vector 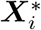 is uniformly bounded.

(A4) There exist positive constants *b* and *B* such that *b* ≤ *λ*_min_(**Σ**_*X*_∗) ≤ *λ*_max_(**Σ**_*X*_∗) ≤ *B*, where **Σ**_*X*_∗ = Var(***X***^∗^), and the information matrix ***I***(***θ***_0_) is positive definite.

(A5) Censoring is independent of the event time given ***X***^∗^, and *P* (*T* ≥ *t*_max_) *>* 0 for some *t*_max_ in the support of the observation time.

#### Remarks.

1. Assumptions (A1) and (A2) require the first-stage estimates 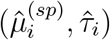 to be uniformly accurate at a rate faster than *n*^−1*/*2^, which allows the plug-in step to be asymptotically negligible. Assumption (A3) prevents instability caused by extreme covariate values and is standard for Cox partial-likelihood asymptotics. Assumption (A4) requires reasonably good behavior of the covariate distribution; in particular, the lower bound *b* requires that *τ*_*i*_ varies across patients, since if *τ*_*i*_ were constant, *β*_2_ would not be identified. Assumption (A5) is the usual independent censoring and risk-set condition of Andersen and Gill (1982).
2. For a twice-differentiable expression surface on a two-dimensional domain, the thin-plate spline attains 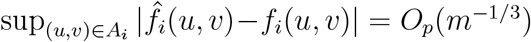 up to logarithmic factors, and 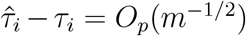 under undersmoothing, so Assumptions (A1) and (A2) reduce to *n*^1*/*2^*m*^−1*/*3^ → 0, i.e. the minimum number of cells per patient growing faster than *n*^3*/*2^. In IMC studies, cell counts per patient are typically in the hundreds to thousands while the number of patients is in the tens to hundreds, so this regime is empirically plausible.

#### Proposition 1

(Oracle property of our plug-in estimator). *Under Assumptions (A1)-(A5), we have* 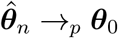 *and*

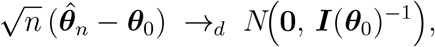

*the same limiting distribution as the infeasible estimator computed with the true* 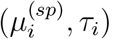; *first-stage estimation is asymptotically negligible*.

*Proof sketch*. Let ***U***_*n*_(***θ***; ***X***) = ∑ _*r*∈*D*_*{****X***_*r*_ − ***S***^(1)^(***θ***, *t*_*r*_; ***X***)*/S*^(0)^(***θ***, *t*_*r*_; ***X***)*}* denote the partial-likelihood score based on covariates ***X*** = (***X***_1_, …, ***X***_*n*_), where 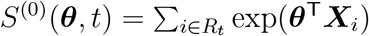 and 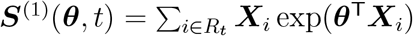. With bounded covariates and *S*^(0)^ bounded away from zero on the observation window, the normalized partial-likelihood score is Lipschitz in the covariates, so 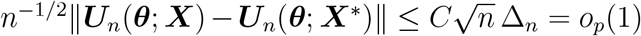 uniformly near ***θ***_0_, for some constant *C*, and also for the observed information. The plug-in criterion thus converges to the same strictly concave limit as the oracle criterion, giving consistency. A Taylor expansion of the score combined with the martingale central limit theorem of Andersen and Gill (1982) then yields the stated limit.

## 3 Simulation Experiments

We conduct simulation studies to compare the proposed method, using spatially adjusted protein summaries with clinical covariates, against existing alternatives. To generate spatially heterogeneous imaging data, cell-level marker intensities measured at spatial coordinates (*u*_*ij*_, *v*_*ij*_). Specifically, for each patient, tumor cell coordinates were sampled from the two-dimensional kernel density estimated from real BC imaging data (Jackson et al., 2020) to preserve spatial information. Figure 2 in the real data application section presents a two-dimensional heatmap and a three-dimensional surface plot of the kernel density for one patient for illustrative purposes. To mimic realistic variations in a protein intensity within the tumor cells, the underlying mean intensity at each coordinate was defined as a mixture of two Gaussian-shaped spatial bumps:

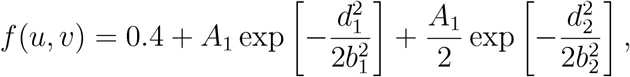

where *d*_*t*_ denotes the Euclidean distance from the *t*-th hotspot center; that is, 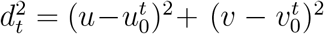, *A*_*t*_ controls the local amplitude with *A*_*t*_ ~ Unif(0.9, 1.6), and (*b*_1_, *b*_2_) determine spatial spread (set to 220 and 180, respectively). The two hotspot centers were randomly drawn as 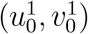 ~ Unif(250, 750) and 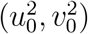 ~ Unif(100, 900) to generate individualized spatial patterns that vary in both location and intensity across patients. Observed cell-level intensities were then obtained as *y*_*ij*_ = *f* (*u*_*ij*_, *v*_*ij*_) + *ϵ*_*ij*_ with 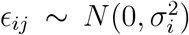, where *σ*_*i*_ ~ Unif(0.1, 0.5). We then simulated three baseline clinical covariates (*z*_1_, *z*_2_, *z*_3_) ~ *N*_3_(0, *I*_3_) for each patient. Patient-level survival times were simulated using an exponential model with rate 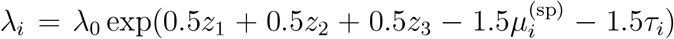, where *λ*_0_ = 0.03, and 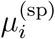 and *τ*_*i*_ denote the true mean spatial PD-L1 intensity and residual spatial variance for *i*-th patient, respectively. Random censoring times were uniformly generated to yield an average censoring rate of approximately 35%.

We implemented our proposed Cox modeling framework using spatially adjusted protein summaries together with clinical covariates. We refer to this model as the Spatially Adjusted Protein Summary (SAPS) model. We compared it with several alternatives: (i) a Cox model that included naïve cell level summaries of protein expression (mean and variance) but ignored spatial information, while adjusting for clinical covariates, (ii) a baseline Cox model with clinical covariates only, and (iii) the geographically weighted (GW) Cox model (Xue et al., 2020). For our proposed method, we set the basis dimension to *k* = 20 for the thin-plate spline GAM term. For the GW method, since clinical covariates did not have cell-level spatial coordinates, we first fitted a Cox model using the clinical covariates to estimate global coefficients. We then computed patient-specific linear predictors, defined as the sum of the linear combinations of clinical covariates weighted by these coefficients, and included them as an offset when fitting the geographically weighted component, which captured spatial variation in protein effects while keeping clinical effects fixed.

We trained the models using two training set of size 40 and 120, each with *m*_*i*_ = 200 tumor cells per patient. We also considered a high cell-count setting where each patient had *m*_*i*_ = 1,000 tumor cells with a training set of size 100. Model performance was evaluated on an independent test set of size 150, using the corresponding *m*_*i*_ values. Predictive performance was evaluated using the Harrell c-index (Harrell et al., 1982, 1984, 1996) and the integrated Brier score (IBS) (Graf et al., 1999). The procedure was repeated 100 times, and we reported the mean and standard deviation of each metric across repetitions. The average computational time in seconds across repetitions was also recorded.

Simulation results with *m*_*i*_ = 200 are presented in Table 1. The proposed method achieved the highest c-index and the lowest IBS. For instance, when *n* = 40, the values were 0.83 and 0.12, respectively. The benefit of using spatial summaries was evident when comparing models with and without them, as the c-index increased from 0.58 (baseline) to 0.83, and the IBS decreased from 0.23 (baseline) to 0.12, with a manageable increase in average runtime. In contrast, the Cox model using naïve cell-level summaries of protein expression had a c-index of 0.78 and IBS of 0.15, yet they were not superior to our proposed method overall. The GW model showed modest discrimination (c-index, 0.69), yet its IBS was nearly twice that of our approach (IBS, 0.23 vs. 0.12). The average runtime of the GW model was roughly similar to that of our proposed method when *n* = 40, but the runtime gap increased as the sample size increased to *n* = 120, with the GW model taking nearly five times longer (22.705 vs. 4.529).

**Table 1:**
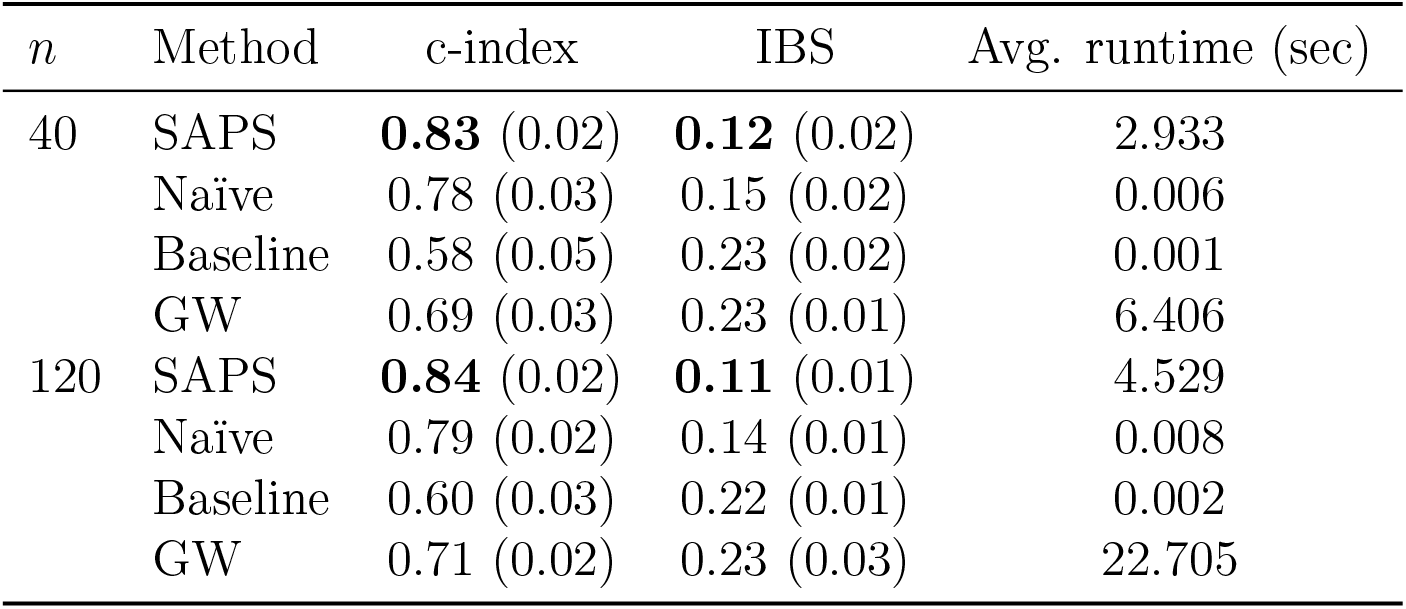
Simulation results when *m*_*i*_ = 200. For each performance metric, the mean is reported with the standard deviation in parentheses. The average computational time in seconds is also provided. The best results are highlighted in boldface.

Table 2 presents the simulation results for the high cell-count setting with *m*_*i*_ = 1,000 tumor cells under *n* = 100. The results were largely consistent with those obtained in the moderate cell-count setting (*m*_*i*_ = 200). Our proposed method achieved the highest c-index and the lowest IBS, indicating strong discrimination and overall predictive accuracy. The other models showed lower discrimination, reduced predictive accuracy, or both. The GW model performed worse than the naïve model in terms of discriminatory ability and also required substantially longer computation time.

**Table 2:** Simulation results when *m*_*i*_ = 1,000 with *n* = 100. For each performance metric, the mean is reported with the standard deviation in parentheses. The average computational time in seconds is also provided. The best results are highlighted in boldface.

| Method | c-index | IBS | Avg. runtime (sec) |
| --- | --- | --- | --- |
| SAPS | **0.85** (0.02) | **0.11** (0.02) | 14.672 |
| Naïve | 0.79 (0.02) | 0.13 (0.02) | 0.012 |
| Baseline | 0.60 (0.03) | 0.22 (0.01) | 0.002 |
| GW | 0.71 (0.03) | 0.23 (0.01) | 18.384 |

We additionally compared the plug-in (SAPS) and oracle Cox estimates of *β*_1_ and *β*_2_, fit to the same simulated outcomes with *n* = 100 fixed and *m*_*i*_ ∈ *{*50, 100, 200, 500, 1,000*}*, over *B* = 100 replications. Figure 1 shows the paired difference between the two estimates by *m*_*i*_. The plug-in estimates are attenuated toward zero relative to the oracle at small *m*_*i*_, and this gap narrows as *m*_*i*_ increases, supporting the claim that first-stage estimation becomes asymptotically negligible.

**Figure 1:**
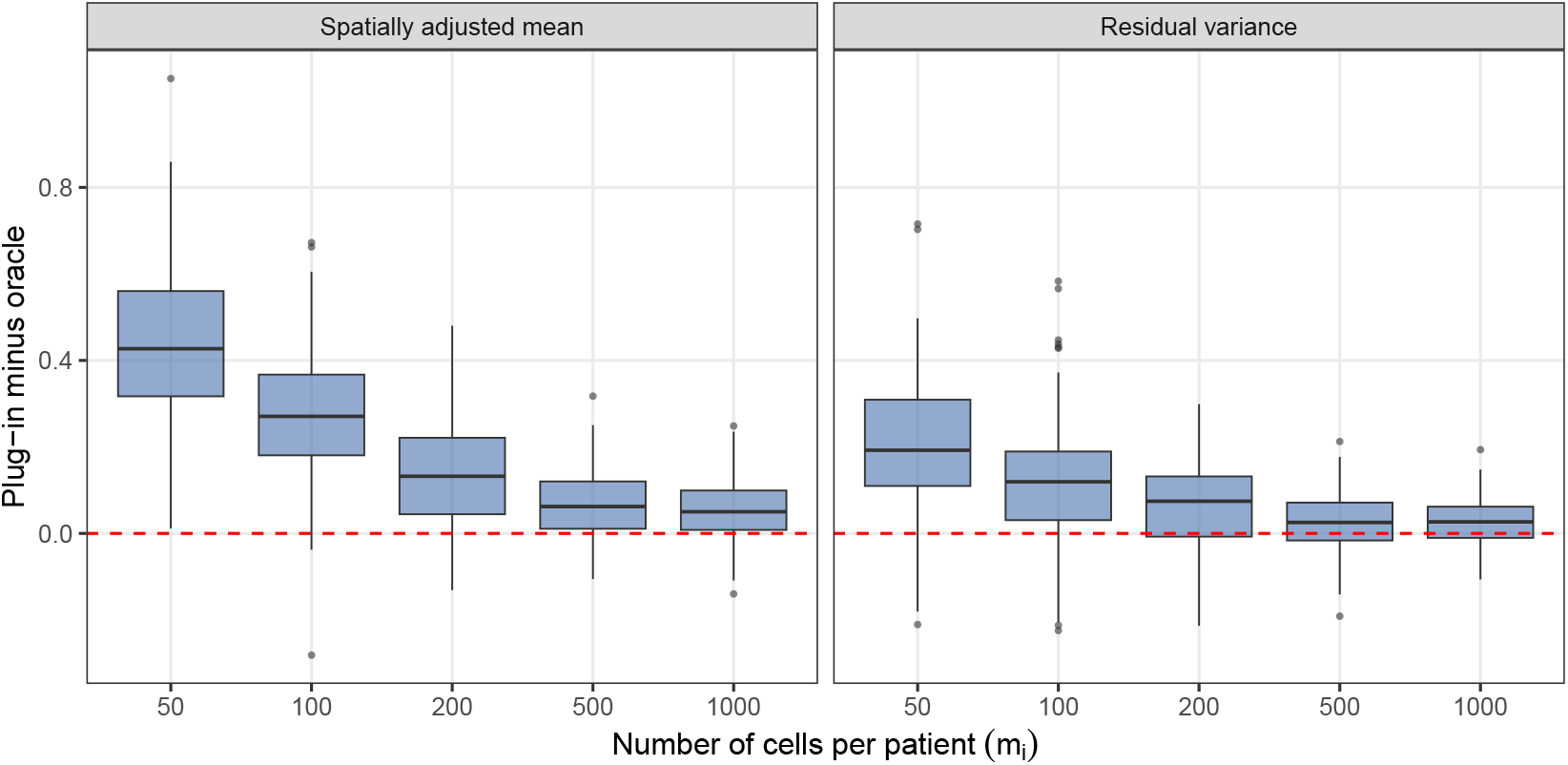
Difference between the plug-in (SAPS) and oracle Cox coefficient estimates (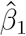 for the spatially adjusted mean and 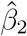 for the residual variance), by number of cells per patient *m*_*i*_, paired within each of *B* = 100 replications with *n* = 100 patients. The dashed line indicates zero difference.

**Figure 2:**
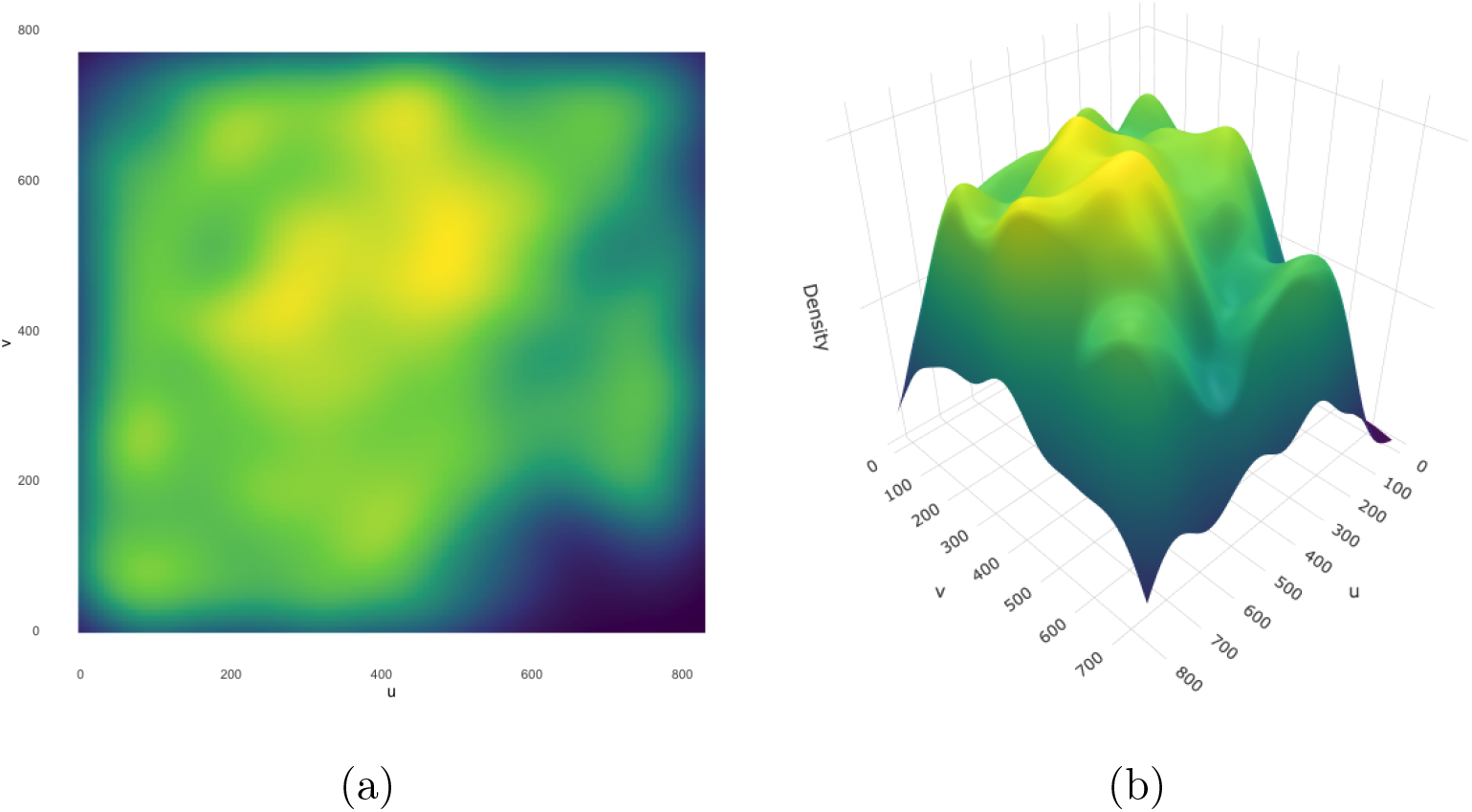
Kernel density estimates of tumor cell coordinates (*u, v*) for patient ID 97 in the BC cohort (Jackson et al., 2020): (a) two-dimensional heatmap; (b) three-dimensional surface plot of the kernel density.

## 4 Real Data Application: Jackson et al. (2020) Study

We applied our proposed method to imaging mass cytometry (IMC) data obtained from tumor tissue of patients with BC, specifically from a spatially resolved single-cell pathology study (Jackson et al., 2020). The dataset included 285,851 single cells from 100 patients, with each cell containing spatial coordinates and CD3E marker intensity measurements. CD3E is a component of the T-cell receptor-CD3 complex and plays an essential role in T-cell activation and adaptive immune response, as described in the Human Protein Atlas database (Uhlén et al., 2020). The number of cells per patient varied, with a median of 2,801 cells (IQR: 2,000 to 3,444). We excluded 9 cases of HR-HER2+ disease and 1 case with a missing subtype from the analysis. The final dataset comprised 90 patients, 25 of whom died, with a median overall survival time of 191 months and subtypes classified as either HR-positive tumors or triple-negative breast cancer (TNBC). This approach was adopted because HR-positive tumors share core biological features and are often analyzed together in survival and prognostic studies when sample sizes are limited. In contrast, TNBC represents the most biologically and therapeutically distinct subtype, characterized by the absence of endocrine and HER2 targets, higher immunogenicity, and distinct immune microenvironment patterns (Zagami & Carey, 2022). Patient-level clinical information included age at diagnosis, subtype, overall survival time, and event status.

The proposed method and other competing methods, including naïve, baseline, and GW models, were applied as described in Section 3. The 5-fold cross-validation was used to fit each model on the training set and evaluate performance metrics on the test set, with the final measures averaged across the folds. Table 3 reports performance metrics for the proposed method and alternative approaches, reported as means across cross-validation folds. Our proposed method achieved the highest c-index and the lowest IBS, indicating superior predictive performance. The naïve and baseline models had IBS values similar to our model, but their c-index values were lower. In particular, the Cox model with naïve cell-level summaries of protein expression did not improve discrimination between high- and low-risk BC patients, as reflected by only a marginal increase in the c-index from 0.68 in the baseline model to 0.69 in the naïve model. In addition, the GW model showed both the lowest c-index and the highest IBS among all methods. These findings highlight the practical advantage of combining spatially adjusted protein summaries with clinical covariates for improved survival prediction in BC patients when the data consist of cell-level marker intensities measured at spatial coordinates.

**Table 3:** Analysis results from a spatially resolved single-cell pathology BC study, reported as means across cross-validation folds. The best results are highlighted in bold.

|  | SAPS | Naïve | Baseline | GW |
| --- | --- | --- | --- | --- |
| c-index | **0.73** | 0.69 | 0.68 | 0.61 |
| IBS | **0.16** | 0.17 | 0.17 | 0.24 |

## 5 Discussion

A major motivation for SAPS was the lack of survival models that adequately account for spatial structure in spatial proteomics data. To address this, protein expression is summarized at the cell level using a GAM-based approach that captures spatial patterns (i.e., intensity). For each patient, two spatial summaries, the spatially adjusted mean and residual variance, are then estimated (see Section 2.1 for details) and regressed in a Cox model to evaluate their association with survival with covariate adjustment. This distinguishes SAPS from TopKAT, a recently developed method for spatial proteomics data that performs a global association test for topological structure in the spatial distribution of cells and sample-level outcomes within a Cox modeling framework. In contrast, SAPS estimates patient-level spatial summaries and models them as covariates in a Cox model with adjustment for clinical variables. This leads to patient-level quantities that can be related to survival risk, whereas TopKAT focuses on a global test statistic. The regression-based formulation of SAPS also makes it straightforward to incorporate clinical covariates and supports survival prediction, while TopKAT is primarily oriented toward hypothesis testing.

Recent studies in triple-negative breast cancer (TNBC) have shown that tumor microenvironments are characterized by risk markers such as Ki67+ (*p* = 0.052) and vimentin+ (*p <* 0.01) tumor cells, which correlate with poor survival (Foroughi Pour et al., 2026). Building on our analyses of BC patients using the CD3 epsilon (CD3E) marker, the SAPS-derived spatially adjusted summaries could help stratify patients for immunotherapy in TNBC, where spatial T-cell patterns are established predictors of response, even in tumors with variable or low PD-L1 expression (Hammerl et al., 2021). The improved predictive performance of our method supports its future integration into routine pathology workflows. Extensions include multi-marker panels (e.g., combining CD3E with PD-L1, CD8, or cancer-associated fibroblast markers), integration with spatial transcriptomics for multi-omics neighborhood analysis, or application to pre-treatment biopsies in the neoadjuvant setting. Validation in larger, treatment-annotated BC cohorts that include racial diversity will further enhance generalizability.

While SAPS has demonstrated promise in improving survival prediction through simulation studies and real data applications, several limitations should be acknowledged. First, our method focuses on cell-level summarization of protein expression for a given protein known to be associated with survival. However, integrating multiple proteins, each measured across thousands of cells, may further enhance predictive performance. Second, the model is built upon the Cox model due to its practicality and popularity; however, its assumptions may be violated, particularly if the proportional hazards assumption does not hold. In future studies, we plan to explore extended models, such as frailty models incorporating random effects or models that do not rely on the proportional hazards assumption, such as the accelerated failure time model.

